# Evidence for a biphasic mechanism of virus transmission of the human metapneumovirus in LLC-MK2 cell monolayers

**DOI:** 10.64898/2026.07.21.739053

**Authors:** Nguyen Huong Tra, Boon Huan Tan, Richard J Sugrue

## Abstract

We examined transmission of the human metapneumovirus (HMPV) in LLC-MK2 cell monolayers using a low multiplicity of infection (moi). In this low moi infection model HMPV transmission initially occurred by localised cell-to-cell transmission, and the virus infectivity remained largely cell associated. At the later stages of infection more widespread virus transmission occurred and was associated with the presence of cell-free virus. The appearance of the cell-free virus correlated with changes in plasma membrane integrity and increased membrane permeability in the cell monolayers. Imaging analysis of HMPV infected cells at the early stages of infection showed the presence of numerous virus filaments attached to the surface of HMPV-infected cells. At the later stages of infection both virus filaments and virus particles with a spherical morphology that was attached to the distal ends of the virus filaments was noted. A proportion of these spherical particles detached from the virus filaments and attached to adjacent non-infected cells at the later stages of infection. The activation of the JNK and MAPKp38 signalling pathways in HMPV-infected cells correlated with increased HMPV replication and appearance of the cell-free virus infectivity. In addition, after the initial phase of STAT1 activation in HMPV-infected cells, both reduced expression of the STAT1 protein and the activated STAT1 protein occurred as the infection proceeded. Collectively, these data provide evidence for a biphasic mode of HMPV transmission involving different virus particle morphologies, a localised virus transmission by virus filaments followed by widespread virus transmission involving cell-free virus particles.

## 1. Introduction

Human metapneumovirus (HMPV) is a negative-sense single-stranded RNA virus of the family *Pneumoviridae*, within the genus *Metapneumovirus*. It is closely related to respiratory syncytial virus (RSV) which is also grouped within the *Pneumoviridae* but in the genus *Orthopneumovirus*. Since its initial identification in Europe [1], HMPV is now recognised as an important virus agent causing respiratory disease in humans and periodic outbreaks of HMPV have been reported in different regions around the world [2, 3]. The HMPV is an important cause of respiratory tract infection in the young and other specific demographic groups, and similar to RSV, the clinical symptoms that are caused by HMPV infections in children range from upper respiratory tract infection to bronchiolitis and pneumonia.

The mature HMPV particle is surrounded by a lipid envelope that is derived from the host cell, into which the virus-expressed fusion (F) and attachment (G) proteins are inserted and are displayed on the surface of the virus envelope. The F protein mediates fusion of the virus and host cell membranes during virus entry [4], while a primary role for the G protein is in mediating virus attachment to susceptible cells prior to cell entry [5]. A single protein complex involving the HMPV F and G proteins has been identified [6], which is similar to protein complex involving the F and G proteins identified for the closely related RSV [7,8]. In the case of RSV, research suggests that this protein complex stabilises the F protein in a prefusion structural conformation [9], and it is similarly presumed that the protein complex involving the HMPV F and G proteins also stabilises the HMPV F protein in its prefusion form. Beneath the virus envelope is a protein layer formed by the virus matrix (M) protein, and a ribonucleoprotein (RNP) complex that is formed by the viral genomic RNA (vRNA), the nucleocapsid (N) protein, the phosphoprotein (P protein), the M2-1 protein and the large (L) protein. Two major HMPV genotypes have been identified based on their genetic characterization which are called HMPV A and B [10–12], although a similar structural organisation of the HMPV particle in both genotypes is presumed.

HMPV morphogenesis occurs on the surface of HMPV-infected cells as long filamentous projections, and evidence for the role for lipid raft microdomains and the cells cortical F-actin network in HMPV morphogenesis has been presented [13]. In this context, F-actin-associated signalling pathways such as Rho GTPases has been demonstrated to play a role in HMPV morphogenesis and transmission [14]. It is established that RSV morphogenesis also involves lipid raft microdomains, the F-actin network and rho GTPase signalling pathways that are associated with F-actin [15], which highlights some of the similarities between HMPV and RSV morphogenesis processes. In this context earlier work on RSV using a permissive cell line and organoid cell system had suggested a biphasic mechanism of virus transmission in permissive cells [16,17]. An early phase involving localised virus transmission mediated by cell-associated virus filaments, which was followed by a later stage involving longer range transmission that is mediated by cell-free infectious virus particles [18]. Since several features of the HMPV morphogenesis process are similar to that reported for RSV, in this current study we have therefore extended our earlier work on HMPV morphogenesis [13] to examine if a similar biphasic mode of HMPV transmission occurs in permissive cells. Understanding the mechanism of HMPV transmission using cell-based models should facilitate the development of antivirus strategies that target the HMPV.

## 2. Materials and Methods

### Cell culture

The cell line LLC-MK2 was purchased (ATTC) and maintained in growth media containing DMEM (Gibco) supplemented with 10% FCS (Gibco) and penicillin and streptomycin (pen/strep) (Gibco) at 37°C in a humidified chamber at 5%CO_2_.

### Virus propagation

The HMPV A2 isolate NCL03-4/174 (HMPV) was obtained from Geoff Toms (University of Newcastle). The HMPV stocks were propagated using LLC-MK2 cells in Maintenance Media (DMEM (Gibco) supplemented with 2.5% (w/v) BSA (Gibco), 1x pen/strep (Gibco) with 0.5 μg/ml trypsin TPCK) at 37°C in a humidified chamber with 5%CO_2_ as described previously [13]. Cells were infected using a multiplicity of infection of 0.1 and once cells showed advanced cytopathic effect (typically at 10 days post-infection) the infected culture media and cells were harvested together, homogenized by gentle agitation with glass beads and 1ml aliquots of the virus preparation were stored at -80°C until required. Prior to use, the HMPV aliquots were flash-thawed, briefly vortexed and then centrifuged (2000rpm x 5 mins) in a tabletop microfuge to remove cell debris.

## 2. Reagents

The anti-F (Mab58) and anti-G (AT1) antibodies have been described previously [19] and were obtained from Geoff Toms, and the anti-M antibody was prepared using recombinant expressed HMPV M protein as described previously [6]. The Evans Blue cell stain (Sigma), anti-mouse IgG conjugated to Alexa Fluor™ 488 were purchased (ThermoFisher). The following primary antibodies were purchased; anti-STAT1 (BD Biosciences), anti-pSTAT1(BD Biosciences), anti-JNK (Cell signaling technology), anti-pJNK (Cell signaling) technology, p38MAPK (Cell signaling technology), pp38MAPK (Cell signaling technology), and anti-actin (Sigma). The lactose dehydrogenase levels in the tissue culture media were measured using the Lactose Dehydrogenase assay (Roche) following the manufacturer’s instructions.

### Multiple cycle virus infection model

The multiple cycle infections were performed on LLC-MK2 cell monolayers using a multiplicity of infection (moi) of 0.05 or 0.01 as indicated in Maintenance Media. In all cases the cells were incubated at 37°C in a humidified chamber at 5%CO_2_ until the time of harvesting. In this low moi virus infection model no agar overlay media was applied to the infected cell monolayers.

### Separation of cell-free and cell-associated HMPV infectivity

The procedure used to separate cell-free (CF) and cell-associated (CA) HMPV infectivity was based on the procedure previously used to separate cell-free and cell-associated RSV infectivity [16]. After infecting LLC-MK2 cell monolayers using the multiple cycle infection model, at the appropriate time of harvesting the CF-virus was harvested in the overlying tissue culture medium. The CA-virus was harvested by replacing the media that was removed with an equal volume of Maintenance Media into the same well and scraping the cells into it. The virus was released by repeat (twice) rapid freeze-thaw using an ethanol/dry ice bath and then a 37°C water bath. The virus suspension was then briefly vortexed and centrifuged (2000rpm x 5 mins) in a tabletop microfuge to remove cell debris. The CA- and CF-virus preparations were then added to LLC-MK2 cell monolayers in Maintenance Media at 37°C in a humidified chamber at 5%CO_2_ until the time of harvesting.

### Assessing virus titer

This was performed using a modified micro-plaque assay that was based on the microplaque assay used to determine RSV infectivity [20]. The HMPV preparation to be assayed was serially diluted in maintenance media and the diluted inoculums applied to well containing sub-confluent (95% confluency) LLC-MK2 cell monolayers. After a 2 hrs. adsorption period the virus inoculums were removed and replaced with maintenance media and after 48 hrs post-infection the monolayers were fixed with ice cold methanol/acetone (1:1) solution for 10 minutes and then washed extensively with PBS. The cell monolayers were stained using anti-M and anti-mouse IgG conjugated to Alexa Fluor™ 488. The infected cell-stained plaques were imaged using a Nikon Eclipse 80i Microscope (Nikon Corporation, Tokyo, Japan) with an Etiga 2000R camera (Q Imaging, Teledyne Photometrics, Tucson AZ, USA) attached. The infected cells were counted, and it was assumed that each individual infected cell/small cell cluster originated from an infectious virus particle in the virus dilution. Each infected cell detected was referred to as plaque forming unit (pfu) and the results obtained were expressed as pfu/ml.

### Imaging using immunofluorescence microscopy and confocal microscopy

LLC-MK2 cell monolayers on coverslips were infected with HMPV using the multiple cycle infection model. At the appropriate time of infection, the cells were fixed using 4% (w/v) paraformaldehyde (PFA) in PBS and afterwards washed using PBS at 4°C. The PFA-fixed cells were permeabilised using 0.1% (v/v) triton X100 in PBS at 4°C for 15 mins prior to staining. The cells were co-stained using the appropriate primary antibody and Alexa Fluor™ 488 secondary antibody and Evans Blue, and the co-stained cells mounted on glass slides using mounting media (Dakocytomation Fluorescence Mounting Media, Dako, USA). The co-stained cells were examined using a Nikon Eclipse 80i Microscope (Nikon Corporation, Tokyo, Japan) with an Etiga 2000R camera (Q Imaging, Teledyne Photometrics, Tucson AZ, USA) attached. The images of immunofluorescence-stained cells were recorded using Q Capture Pro ver. 5.0.1.26 (Q Imaging, Teledyne Photometrics). For confocal microscopy the stained cells were imaged using a Zeiss LSM 510 laser scanning confocal microscopy using appropriate machine settings with 100x magnification/oil objective lens and images processed using Zen 2.3 software.

### Immunoblotting

The LLC-MK2 cell sheets were washed once with PBS were extracted directly into Laemli buffer (Biorad) and heated at 95°C for 5 minutes. The extracted cell lysates were sonicated briefly using a sonicating probe to reduce sample viscosity and then centrifuged at 13,000 rpm for 2 mins in a table-top microfuge. The proteins were separated by SDS–PAGE, transferred onto PVDF membranes (Immobilon-P, Milipore, USA) and the PVDF membranes were blocked using 5% skim milk (Marvel) in 0.25% Tween 20 in PBS. The membranes were extensively washed in 0.25% Tween 20 in PBS and probed with the respective primary antibody and the appropriate goat anti-mouse or anti-rabbit IgG secondary antibody conjugated to HRP (Sigma). The membranes were finally washed using 0.25% Tween 20 in PBS and the Protein bands were visualised using the ECL system (GE Healthcare, USA). Molecular masses were estimated using Kaleidoscope markers (Biorad, USA).

## 3. Results and discussion

### 3.1 Multi-cycle HMPV infection

In this study we used the HMPV A2 isolate NCL03-4/174 (HMPV) that has been described previously [21]. The LLC-MK2 cell line was used throughout this study since this cell line is permissive for HMPV infection and can be used for HMPV propagation and to investigate HMPV transmission. HMPV-specific antibodies used in this study are anti-F antibody (recognises the HMPV F protein), anti-M (recognises the HMPV M protein) and anti-G (recognises the HMPV G protein). In this study the cells were infected with HMPV using a low multiplicity of infection (moi) and unless otherwise indicated the low moi used was 0.05. This experimental format diluted out any defective interfering particles that may be present in the virus inoculum and enabled us to examine cells infected only with fully infectious virus particles. The infected LLC-MK2 cells were also incubated at 37°C without the application of an agar overlay media to the cell monolayer. Since an agar overlay constrains the distribution of virus released from the cell i.e. cell-free virus infectivity, the absence of the agar overlay also allowed us to distinguish localised virus transmission and long-range virus transmission across the overlay by cell-free virus infectivity

LLC-MK2 cell monolayers were mock-infected and infected with HMPV and at 3, 7 and 10 days post-infection (dpi) the cells were fixed and co-stained using Evans Blue (EB) and each of the HMPV-specific antibodies (Fig. 1). The EB is a non-specific cell stain that allows imaging of all the cells in the monolayer in the same field of view. The co-stained cells were imaged using immunofluorescence (IF) microscopy to detect antibody-stained cells and the EB-stained cell monolayer. Mock-infected cells showed the absence of staining with each of the virus-specific antibodies, and the presence of the intact EB-stained continuous cell monolayer (Fig. 1A(i), Fig. 1B(i) and 1C(i)). At 3 dpi staining of the infected cell monolayers with each virus-specific antibody revealed only individual stained cells consistent with the presence of individual infected cells within the cell monolayer (Fig. 1A(ii), Fig. 1B(ii) and 1C(ii)). No background staining was noted at 3 dpi with each of the virus-specific antibodies when examined by IF microscopy indicating that any background staining was below the level of detection by IF microscopy. At 7dpi we noted the appearance of larger antibody-stained cell clusters consisting of up to 5 cells (Fig. 1A(iii), Fig. 1B(iii) and Fig. 1C(iii)), which were similar in appearance to the presence of HMPV-induced cell syncytia. By 10 dpi we noted that these infected cell clusters increased in size, and the presence of smaller cell clusters together with individual infected cells were also noted. (Fig. 1A(iv), Fig. 1B(iv) and 1C(iv)). At 10 dpi the larger infected cell monolayers showed extensive cell syncytia-like formations. These data obtained using the low moi infection model were consistent with localised cell-to-cell spread of HMPV infection from within the cell monolayer to generate the larger infected cell clusters and syncytia at the later times of infection.

**Figure 1.**
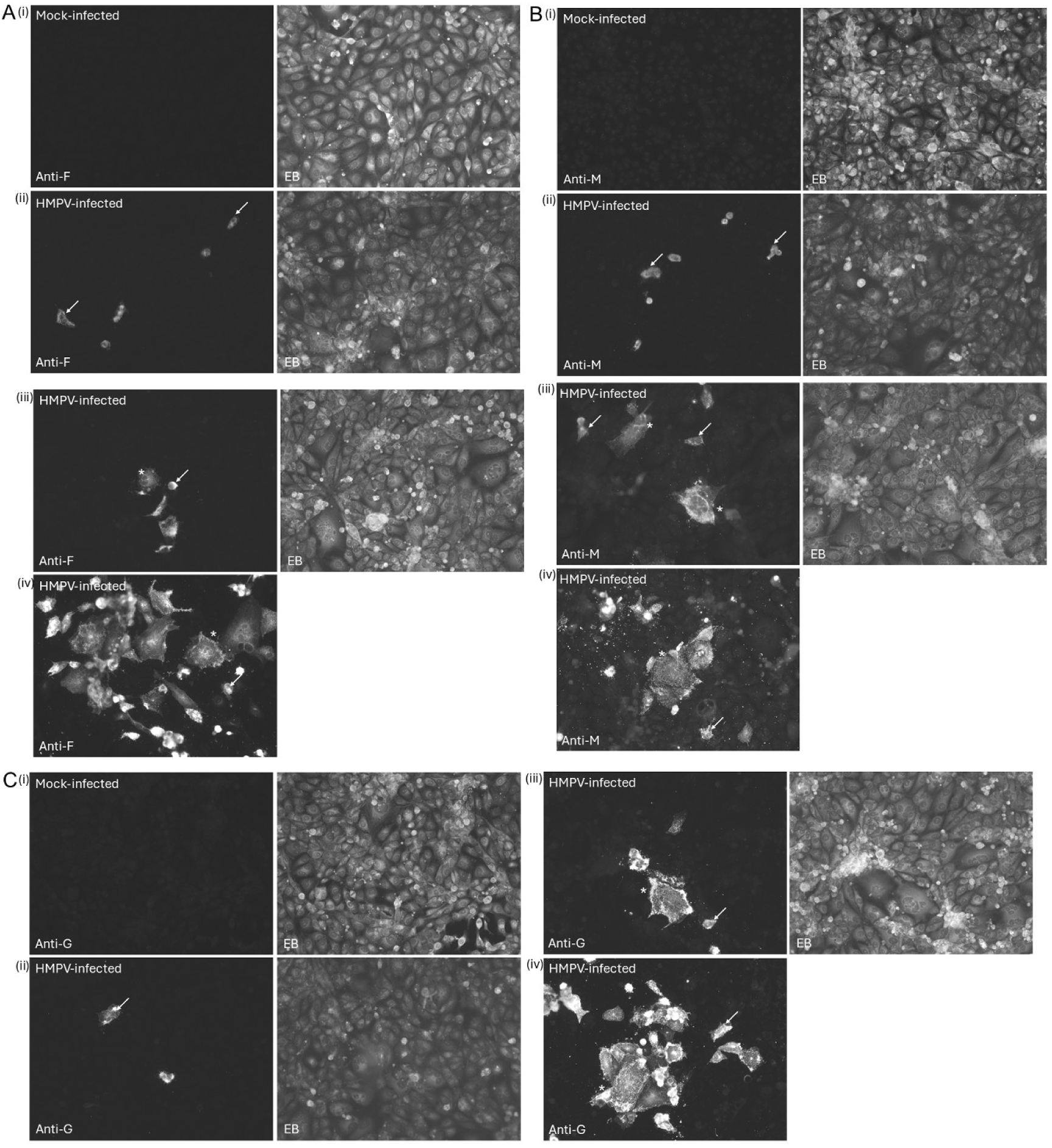
Multiple cycle infection of HMPV in LLC-MK2 cell monolayers. Cell monolayers were (i) Mock-infected and (ii to iv) infected with HMPV (HMPV-infected) as indicated using a moi of 0.05. At (ii) 3 day-post infection (dpi), (iii) 7 dpi, and (iv) 10dpi the monolayers were co-stained with Evens Blue (EB) and **(A)** anti-F, **(B)** anti-M and **(C)** anti-G, as indicated. Also shown are the (i) mock-infected cells co-stained with each individual virus-specific antibody and EB. The co-stained cells were imaged by immunofluorescence microscopy (objective x20 Magnification). The individual infected cells (white arrows) and clusters of infected cells (*) are highlighted.

We examined the multiple-cycle HMPV model to determine the relative levels of cell-associated and cell-free virus infectivity. LLC-MK2 cell monolayers were infected with HMPV and at between 1 and 10 dpi the relative amounts of the cell-associated (CA) and cell-free (CF) virus infectivity were measured (Fig. 2A). The infectivity assays were performed at 2-day intervals within this time interval i.e. at 1, 3, 5, 7 and finally at 10 dpi. We presumed that at 1 dpi any infectivity recorded may arise largely from input virus that was used to carry out the infection. However, after 3 dpi there was a noticeable increase in the CA-virus infectivity, that continued to increase as the infection proceeded. While there was a lag period before significant levels of CF-infectivity could be detected, a significant increase in CF-virus infectivity occurred only after 7 dpi. However, throughout the time period of the multiple cycle infection the CF-infectivity remained significantly lower that the level of the CA-virus infectivity measured, and even at 10 dpi when there was extensive CPE the CF-infectivity was approximately 50% of the CA-virus infectivity levels. Fresh LLC-MK2 cell monolayers were challenged with equal volumes of each of the CA-virus and CF-virus preparations obtained at 1, 3, 5, 7 and 10 dpi and these were stained with anti-M and imaged using IF microscopy (Fig. 2B). Low levels of virus infectivity in the LLC-MK2 cell monolayers were detected at 3 dpi when challenged with both preparations. However, at between 5 and 10 dpi larger numbers of antibody-stained infected cells was observed in cells challenged with the CA-virus preparation compared with cells challenged with the CF-virus preparation. Collectively, these data indicated that the bulk of the virus infectivity remained largely cell-associated even at the later stages of infection, but that significant levels of cell free-virus was detected only at the later stages in the multiple cycle infection.

**Figure 2.**
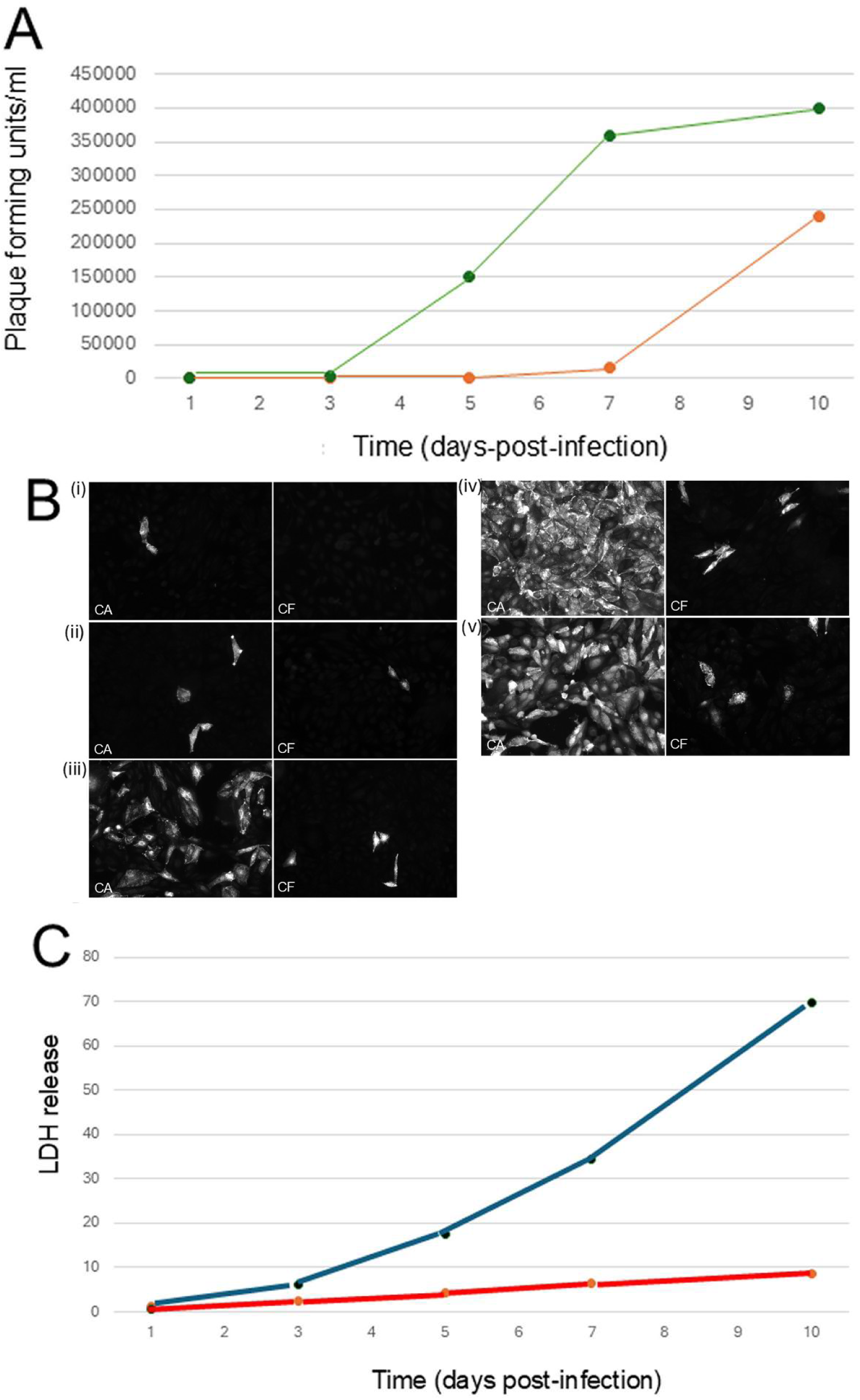
The presence of cell-free virus infectivity correlates with change in cell membrane integrity. LLC-MK2 cell monolayers were infected with HMPV using a moi of 0.01 and **(A)** at between 1 day-post infection (dpi) and 10 dpi the cell-associated (green line) and cell-free (orange line) virus infectivity were estimated. The average infectious titres from duplicate measurements at each time point are shown (SE ±5% for each data point). **(B)** Equal volumes of the cell-associated (CA) and cell-free (CF) samples harvested at (i) 1dpi, (ii) 3 dpi, (iii) 5 dpi (iv) 7 dpi and (v) 10 dpi were used to infect fresh LLC-MK2 cell monolayers and at 24 hpi the cells were stained with anti-M and imaged by immunofluorescence microscopy (objective x20 magnification). **(C)** At between 1 dpi and 10 dpi the corresponding levels of lactose dehydrogenase (LDH) in the tissue culture media in both mock-infected (red line) and HMPV infected (blue line) cell monolayers were estimated using the Lactose Dehydrogenase assay (Roche). The LDH release is presented as a percentage of the LDH release measured in detergent permeabilised cells as per the manufacturer’s instructions. The average infectious titres from duplicate measurements at each time point are shown (SE ±5% for each data point).

In a parallel analysis we also measured Lactate dehydrogenase (LDH) release from the LLC-MK2 cells to assess the membrane permeability of the cells during HMPV infection in the multiple cycle infection model (Fig. 2C). When cells are damaged the change in membrane integrity leads to the release of LDH into the extracellular environment, where its activity can be detected and quantified using the LDH assay. We had previously used this approach to examine membrane permeability changes in RSV infection using a similar multicycle infection model [16]. In this analysis the LDH levels released from mock-infected and HMPV-infected cells were recorded and compared with the LDH levels in a positive control where total LDH levels were released from the monolayer following detergent treatment. The LLC-MK2 cell monolayers were either mock-infected or infected with HMPV and at between 1 and 10 dpi the LDH levels in the tissue culture media was measured and presented as a percentage of the positive control. In the mock-infected condition there were low LDH levels recorded throughout the duration of the infection. At the early stage this accounted for less than 2%, and by 10 dpi (i.e. end point of the assay) an 8% increase was recorded, the observed increase at 10 dpi was presumably a consequence of natural physiological processes at this extended time of cell culturing. However, in the HMPV infection cell monolayer there was a significant increase in LDH levels after 5dpi, and by 10 dpi this accounted for 70% of the positive control. This indicated an increase in membrane permeability during HMPV infection, and that this increase in cell membrane permeability occurred at a time just prior to the appearance of the CF-virus infectivity. This suggested that the appearance of the cell-free virus particles was associated with changes in the membrane integrity in these cells. These observations were generally similar to the observations made using an RSV using a similar multiple cycle infection model [16] where the appearance of the cell-free virus infectivity was associated with changes in the cell membrane permeability. However, it was noteworthy that the time taken for HMPV-infected cells to show syncytia was significantly longer than in similar permissive cells infected with RSV using a multiple cycle infection. The reason for this is currently unclear, but it may be related to the differences in the physiology of the different cells used to culture the viruses and differences in the biological properties of the HMPV and RSV isolates at low moi.

### 3.2 Different HMPV particle morphologies are detected using the multicycle infection model

Analysis of the virus particle morphology at 3 and 10 dpi was performed using IF microscopy at higher magnification. Since the distribution of the HMPV glycoproteins that cover the virus particles would more clearly define the virus particle morphology this comparison was performed on individual anti-F and anti-G-stained HMPV-infected cells at 3 and 10 dpi (Fig. 3). At 3 dpi the anti-F and anti-G-stained cells exhibited a predominantly filamentous staining pattern that was consistent with the presence of virus filaments. However, at 10 dpi we also noted the presence of an addition straining pattern on both anti-F and anti-G-stained cells that appeared as brightly stained spots, suggesting an additional particle morphology at the later time of infection.

**Figure 3.**
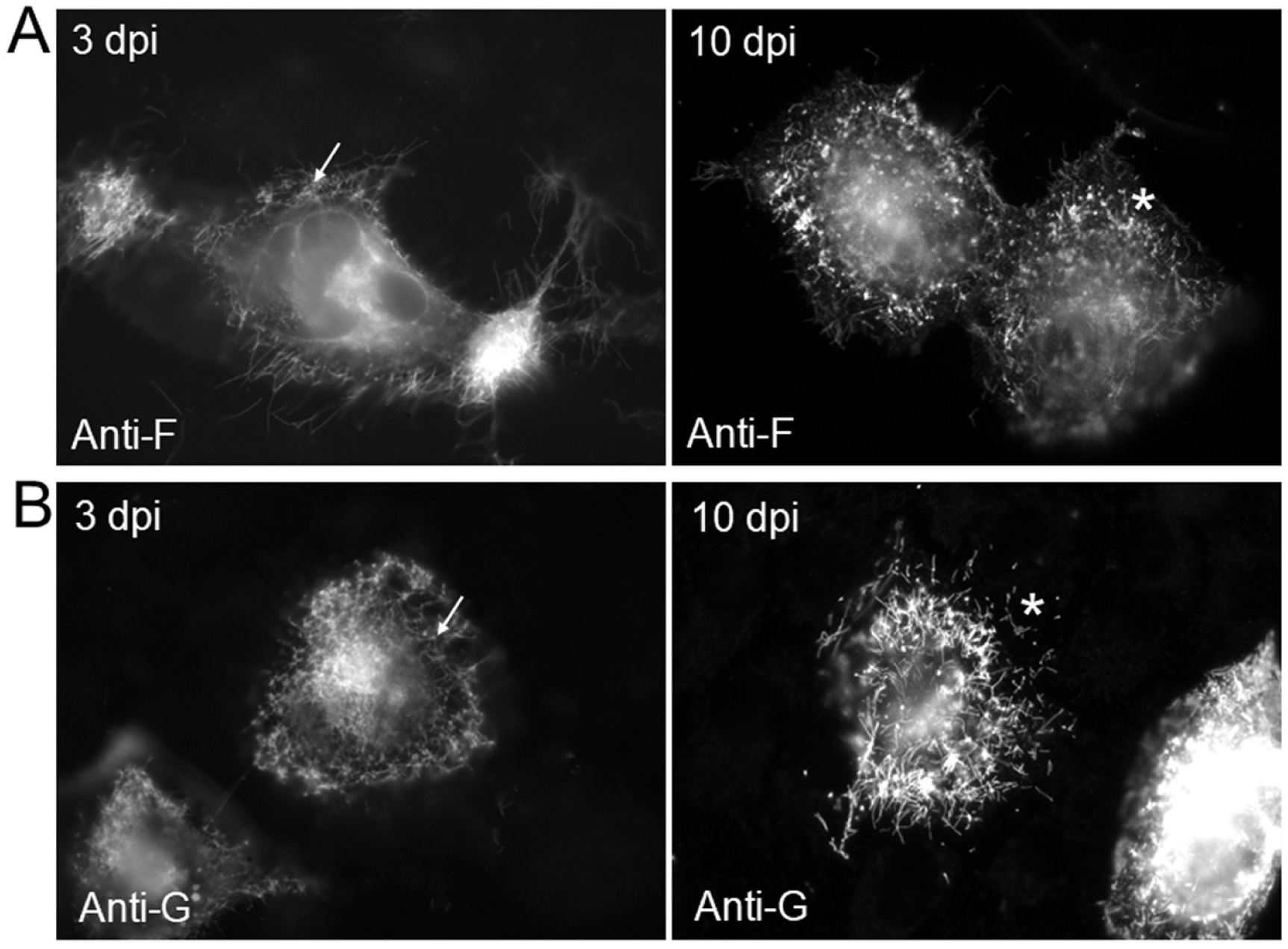
Comparison of anti-F and anti-G stained HMPV-infected cells at 3 and 10 days post-infection using immunofluorescence microscopy. LLC-MK2 cell monolayers were infected with HMPV using a moi of 0.05. and at 3 days postinfection (3dpi) and 10 days post-infection (10 dpi) the cells were stained with **(A)** anti-F and **(B)** anti-G and the cells imaged using immunofluorescence microscopy (objective x100 (oil) magnification). The virus filaments (white arrows) and regions on the infected cell exhibiting the alternative spotted staining pattern (*) are highlighted.

The cell monolayers were examined in greater detail using confocal microscopy to image the virus particle morphology that form on the cell monolayer at 10 dpi. The anti-F and EB-co-stained cells were imaged at a focal plane that allowed imaging of the cell periphery (Fig. 4A) and near the cell top (Fig. 4B). In this image analysis the virus filaments can be seen to be extending from the surface of the infected cells to the adjacent non-infected EB-stained cells. A similar analysis was performed on anti-M and EB-co-stained cells using confocal microscopy to image the cell periphery (Fig. 4C) and near the cell top (Fig. 4D). This showed similar virus filaments extending from the surface of the infected cells to the adjacent non-infected EB-stained cells. A more detailed analysis of the surface of anti-F and EB-co-stained infected cells was performed using confocal microscopy (Fig. 5A). This showed the virus filaments on the cell surface, and in addition the appearance of an anti-F-stained structure with a more spherical morphology at the distal ends of a proportion of the virus filament in the field of view (Fig. 5B(i)). In this report we refer to these features as virus particles with rounded morphology (VPRM) to distinguish them from the filamentous morphology adopted by the virus filaments. In addition, we also noted the appearance of the anti-F-stained spherical structures on the surface of the adjacent non-infected cells which resembled the VPRM that formed at the distal ends of the virus filaments but that had completely detached from the virus filaments (Fig. 5B(ii)). A densitometry analysis was used to measure anti-F antibody staining intensity along the length of the virus filaments using a representative virus filament (Fig. 5C(i)). The increased staining intensity at the distal end of the virus filament (Fig. 5C(ii) and (iii)) suggested an accumulation of the F protein at the tip of the virus filaments within a district structure that forms on the virus filaments.

**Figure 4.**
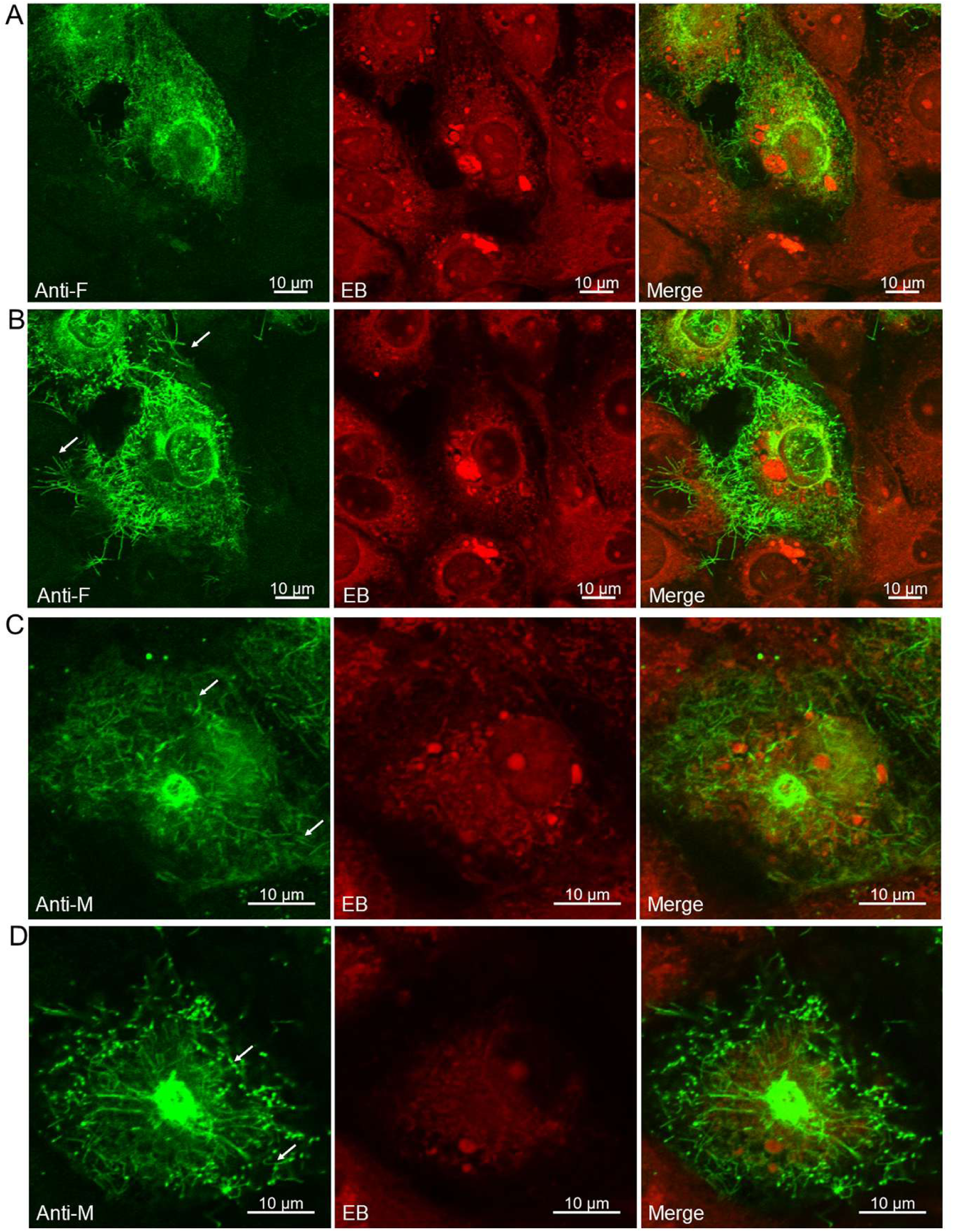
Imaging of HMPV-infected cells using confocal microscopy. LLC-MK2 cell monolayers were infected with HMPV using a moi of 0.05. At 10 day-post infection the cells were co-stained with **(A and B)** anti-F and **(C and D)** anti-M and Evans Blue (EB) as indicated. The cells were examined by confocal microscopy to highlight the (**A and C)** cell periphery and **(B and D)** the cell top. The virus filaments (white arrows) are highlighted. Each image channel together with Merge image channel are presented.

**Figure 5.**
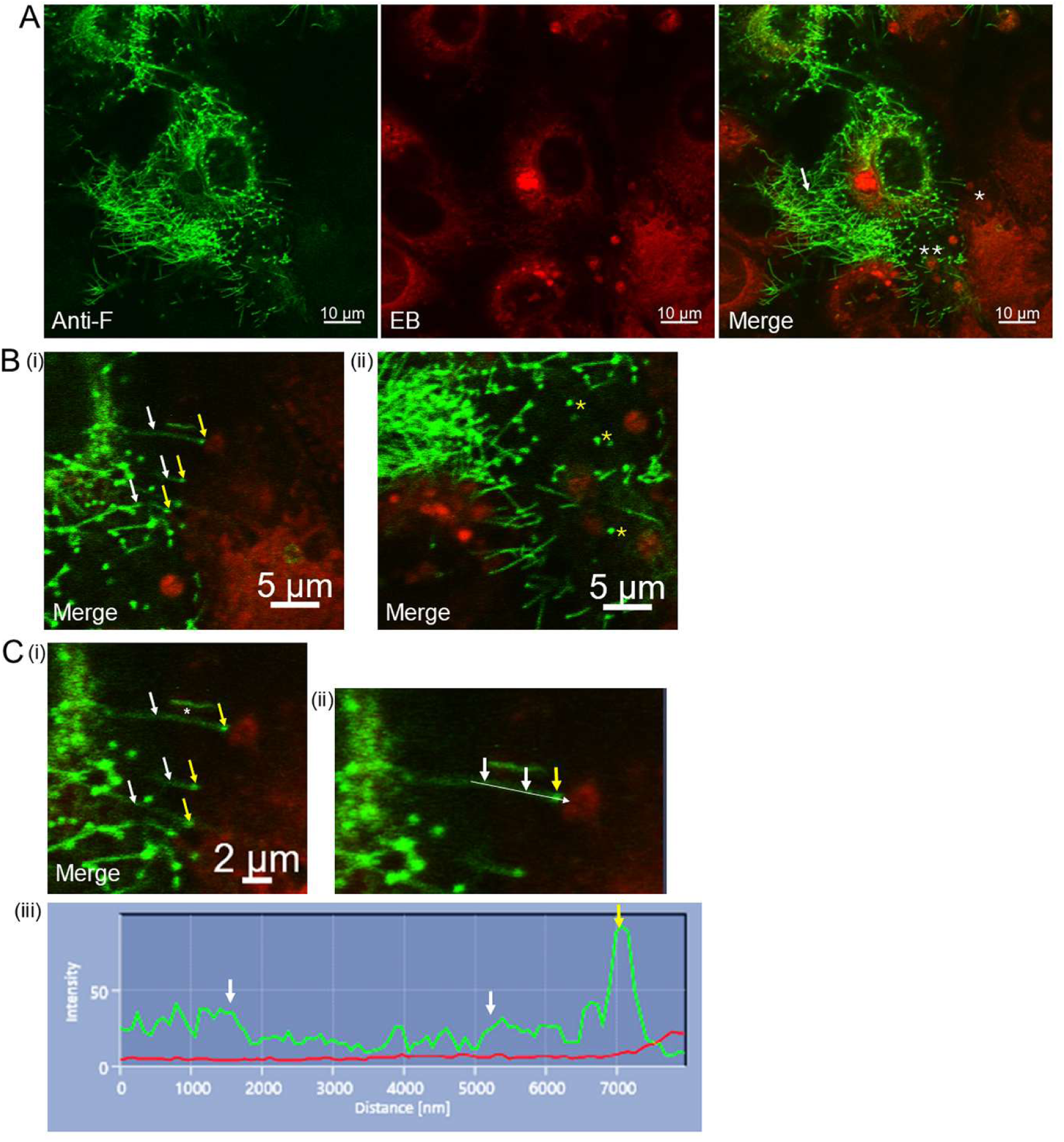
Imaging of anti-F stained HMPV-infected cells using confocal microscopy reveals two virus particle morphologies. LLC-MK2 cell monolayers were infected with HMPV using a moi of 0.05. At 10 day-post infection the cells were co-stained with anti-F and Evans Blue (EB) as indicated. **(A)** The cells were examined by confocal microscopy to highlight the cell top. The virus filaments (white arrows) are highlighted, and each image channel and the Merge image channel are presented. **(B)** (i) and (ii) are enlarged images taken from plate **(A)** at the region highlighted by (*) and (**) respectively. (i) the virus filaments (white arrows) and (ii) the increased anti-F staining at the distal ends of the virus filaments (yellow arrows) are highlighted. (ii) The spherical structures that are detached from the virus filaments that are attached to the surface of the adjacent non-infected cells (yellow *) are highlighted. Only the Merge image is presented. **(C)** Quantification of anti-F staining in the virus filaments was performed using densitometry (i) The virus filaments (white arrows) and the increased anti-F staining at the distal ends of the virus filaments (yellow arrows) are highlighted. (ii) The virus filament highlighted (*) in (i) was examined by densitometry and only the Merge image is presented. The white line highlights the region of the image examined and (iii) is the resulting densitometry scan. The green line represents the anti-F antibody staining intensity, and the red line represents the EB staining intensity. The white arrows correspond to the image intensity at regions along the virus filament, and the yellow arrow corresponds to the image intensity at the distal end of the virus filament that are indicated by the corresponding arrows in (ii).

A similar imaging analysis was also performed on anti-M and EB-co-stained cells using confocal microscopy (Fig. 6A), and a similar increased anti-M staining intensity at the distal end of the virus filament was noted that resembled the VPRM (Fig. 6B(i)). In addition, we also noted the presence of the anti-M-stained spherical structures that resembled the VPRM that were not attached to the virus filaments but that were present on the surface of the adjacent non-infected cells (Fig. 6B(i)); that were similar in appearance to the structures detected on the anti-F-stained cells. A similar image analysis was performed using the HMPV-infected cells co-stained with anti-G and EB (Fig. 7A), and the increased anti-G staining intensity at the distal end of the virus filaments that resembled the VPRM (Fig. 7B) was noted. This again suggested an accumulation of the G protein at the distal end of the virus filaments, similar to that observed on virus filaments on cells stained using either anti-F or anti-M. While the F and G proteins are displayed on the surface of virus envelope on the virus particles, the M protein is located beneath the virus envelope. This indicated that the appearance of these VPRM on the distal ends of the virus filaments was not dependant on the virus-specific antibody used in the staining, more so given the different locations of the virus glycoproteins and the M protein in the context of the virus envelope. We estimated the diameter of the VPRM that were attached to the virus filaments on the anti-F (Fig 8A) and anti-M (Fig 8B) stained cells by using densitometry to access the anti-body staining intensity, and we estimated that they had an approximate diameter of between 0.8 to 1μM. Although the signal dispersion from the fluorophore detected in these images obtained by confocal microscopy is likely to give an apparent increase in size of these structures compared with their actual size, the VPRM clearly have a larger width that the virus filaments detected (approximately 400 nm) in the same field of view and under the same labelling conditions.

**Figure 6.**
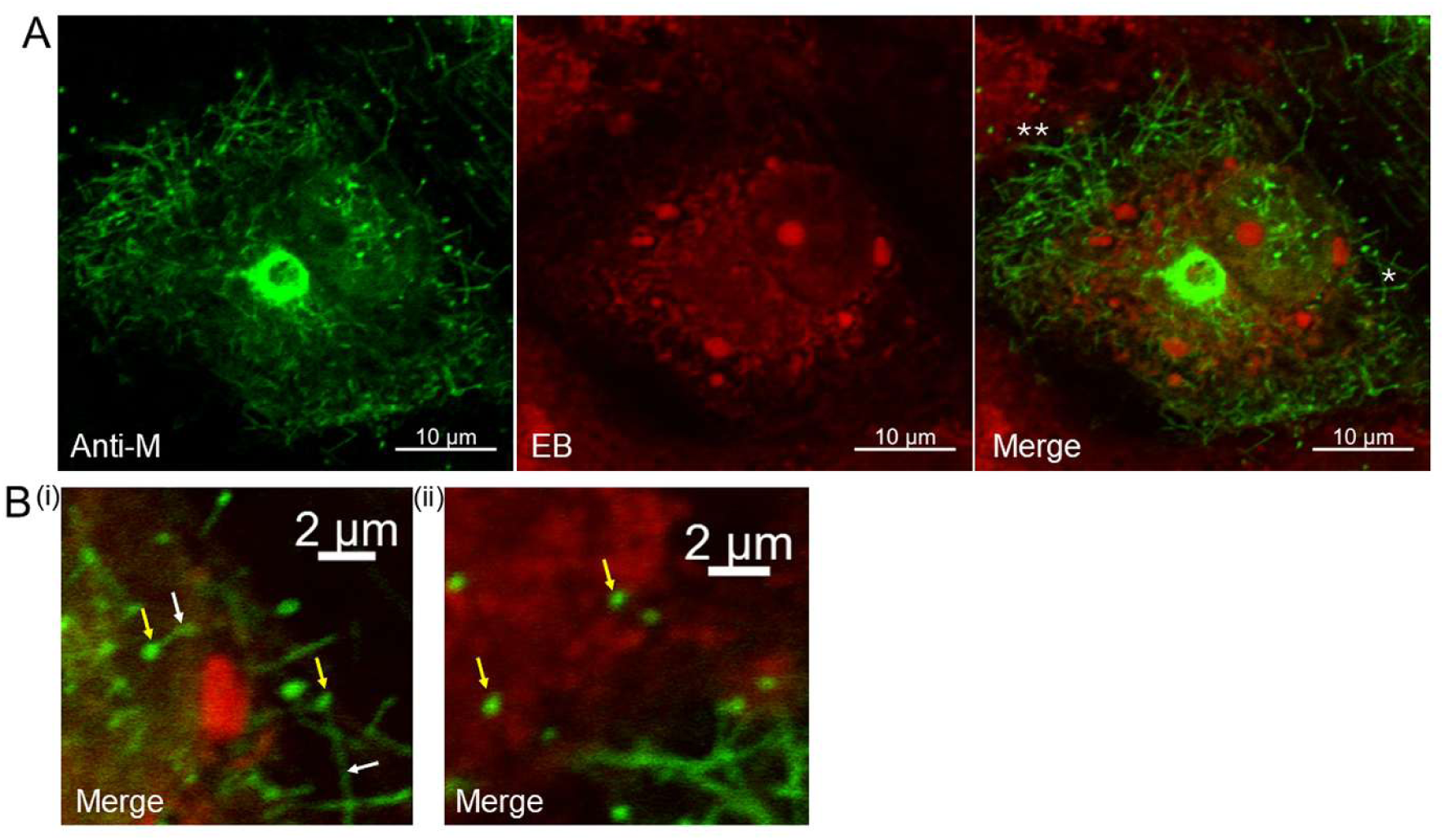
Imaging of anti-M stained HMPV-infected cells using confocal microscopy reveals two particle morphologies. LLC-MK2 cell monolayers were infected with HMPV using a moi of 0.05. At 10 day-post infection the cells were co-stained with anti-M and Evans Blue (EB) as indicated and **(A)** The cells were examined by confocal microscopy to image the virus filaments. Each image channel to together with Merge image channel are presented. **(B)** (i) and (ii) are enlarged images taken from regions in plate **(A)** indicated by (*) and (**) respectively. Only the Merge images are shown. (i) The virus filaments (white arrows) and the spherical structures at the distal ends of the virus filaments (yellow arrows) are highlighted. (ii) The spherical structures that are detached from the virus filaments that are attached to the surface of the adjacent non-infected cells (yellow arrows) are highlighted.

**Figure 7.**
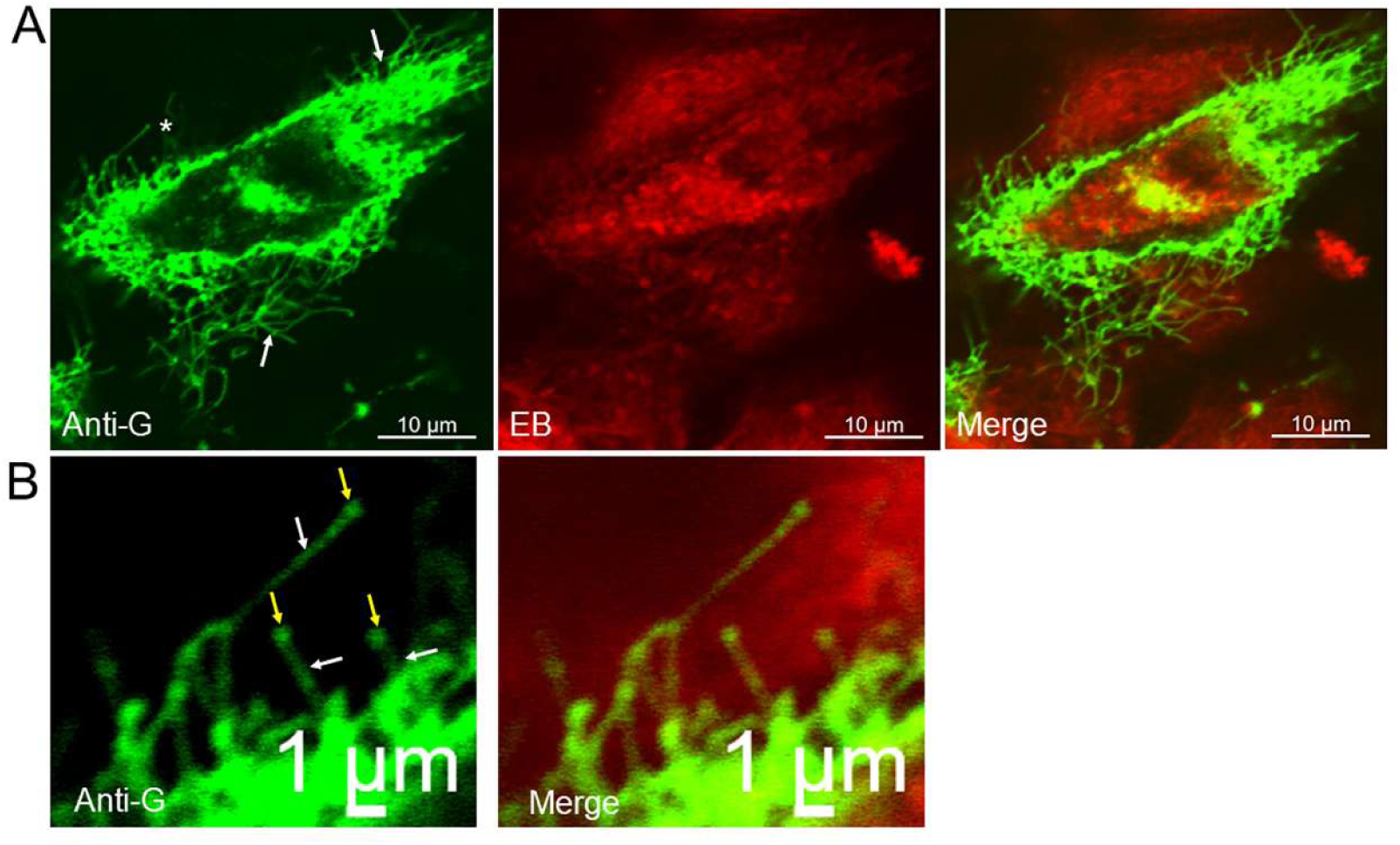
Imaging of anti-G stained HMPV-infected cells using confocal microscopy reveals two particle morphologies. LLC-MK2 cell monolayers were infected with HMPV using a moi of 0.05. **(A)** At 10 day-post infection the cells were co-stained with anti-G and Evans Blue (EB) as indicated. The cells were examined by confocal microscopy to image the virus filaments (white arrows) which are highlighted. Each image channel to together with Merge image channel are presented. **(B)** is an enlarged image taken from the region is highlighted (*) in plate **(A)**. The virus filaments (white arrows) and increased anti-G staining at the distal ends of the virus filaments (yellow arrows) are highlighted. Only the anti-G image and Merge channels are presented.

**Figure 8.**
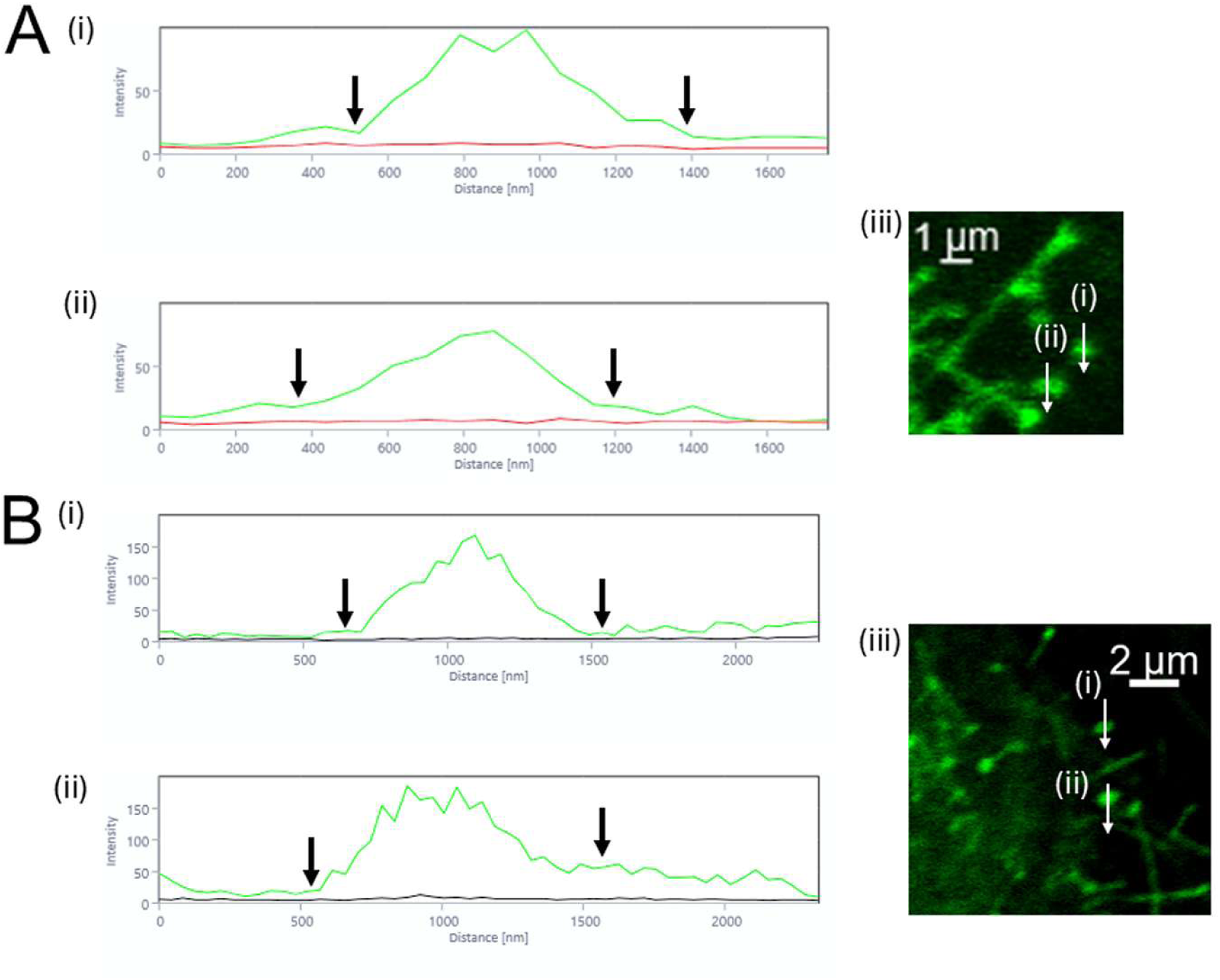
Estimation of the physical dimension of the spherical structures that form on the distal ends of the virus filaments. HMPV-infected LLC-MK2 cells were co-stained with Evans Blue and **(A)** anti-F or **(B)** anti-M. The cells were examined by confocal microscopy to highlight the cell top to image the virus filaments. In each case (i) and (ii) are densitometric scans of the corresponding spherical structures in plate (iii). The white line indicates the region scanned and the arrow indicates the direction of the scan. In the appropriate densitometric scans the green line represents the anti-F and anti-M antibody staining intensity, and the red and black line represents the EB staining intensity. The large black arrows on the densitometric scans demarcates the regions showering increased antibody staining intensity in the scan.

These data suggest that that the virus filaments allowed the spread of the infection of HMPV to neighbouring non-infected cells in the cell monolayer. However, after the virus filaments formed, these virus induced structures gave rise to virus particles with a more spherical morphology that formed at the distal ends of the virus filaments; providing evidence that the virus particles adopted two morphologies. The virus filaments that are attached to the cells are cell-associated virus infectivity, and we presume that once the virus particles with the spherical morphology are detached from the distal ends of the virus filaments, that they form the basis for the cell-free virus infectivity that we detected later in infection.

### 3.3 HMPV particle formation and transmission is associated with activation of the JNK and MAPKp38 signalling pathways

In a final analysis we examined the effect of the virus infection on the activation of the STAT1, JNK and MAPKp38 signalling pathways in the cell monolayers. The activation of the STAT1 protein plays a role in interferon signalling [22] and could potentially create an antiviral response in the HMPV-infected cell monolayer. In addition, both JNK and MAPKp38 signalling pathways have been proposed to facilitate virus replication and production in virus-infected cells in the closely related RSV [23,24]. LLC-MK2 cell monolayers were mock-infected and infected with HMPV and at between 1 and 6 dpi the cells were extracted with boiling mix and analysed by using immunoblotting. This was performed using both the anti-G and anti-M virus-specific antibodies (Fig. 9A) and using antibodies that recognise the STAT1, JNK and MAPKp38 proteins and the activated forms of each protein i.e. pSTAT1, pJNK and pMAPKp38 proteins (Fig. 9B). At between 1 and 6 dpi there was a gradual increase in the G protein and M protein levels (Fig. 9A) that confirmed the HMPV infection and was consistent with increased virus replication over the time of infection. The absence of these protein bands in the mock-infected cell lysates confirmed the specificity of anti-G and anti-M antibodies. The analysis of the STAT1, JNK and MAPKp38 signalling pathways was also examined using the same cell lysates (Fig. 9B). While the presence of the pSTAT1 protein was detected at 1 and 2 dpi, there was a concomitant decrease in both STAT1 protein levels and pSTAT1 levels at between 2 and 6 dpi. While there was no decrease in JNK and MAPKp38 protein levels at between 1 and 6 dpi, the presence of activated forms of these proteins (i.e. pJNK and pMAPKp38) was detected at 2 dpi. Although the levels of pJNK and pMAPKp38 appeared to peak at 4 dpi, low levels of pJNK and pMAPKp38 could still be detected up to 6 dpi. The detection of pJNK and pMAPKp38 appeared at a time that coincided with the increased levels of HMPV infectivity in the multiple cycle virus infection model, and just prior to increases in the levels of CF-infectivity. However, it is currently unclear if the activation of the JNK and MAPKp38 signaling pathways mediate the release of the cell-free virus infectivity that is detected at the later stages of infection.

**Figure 9.**
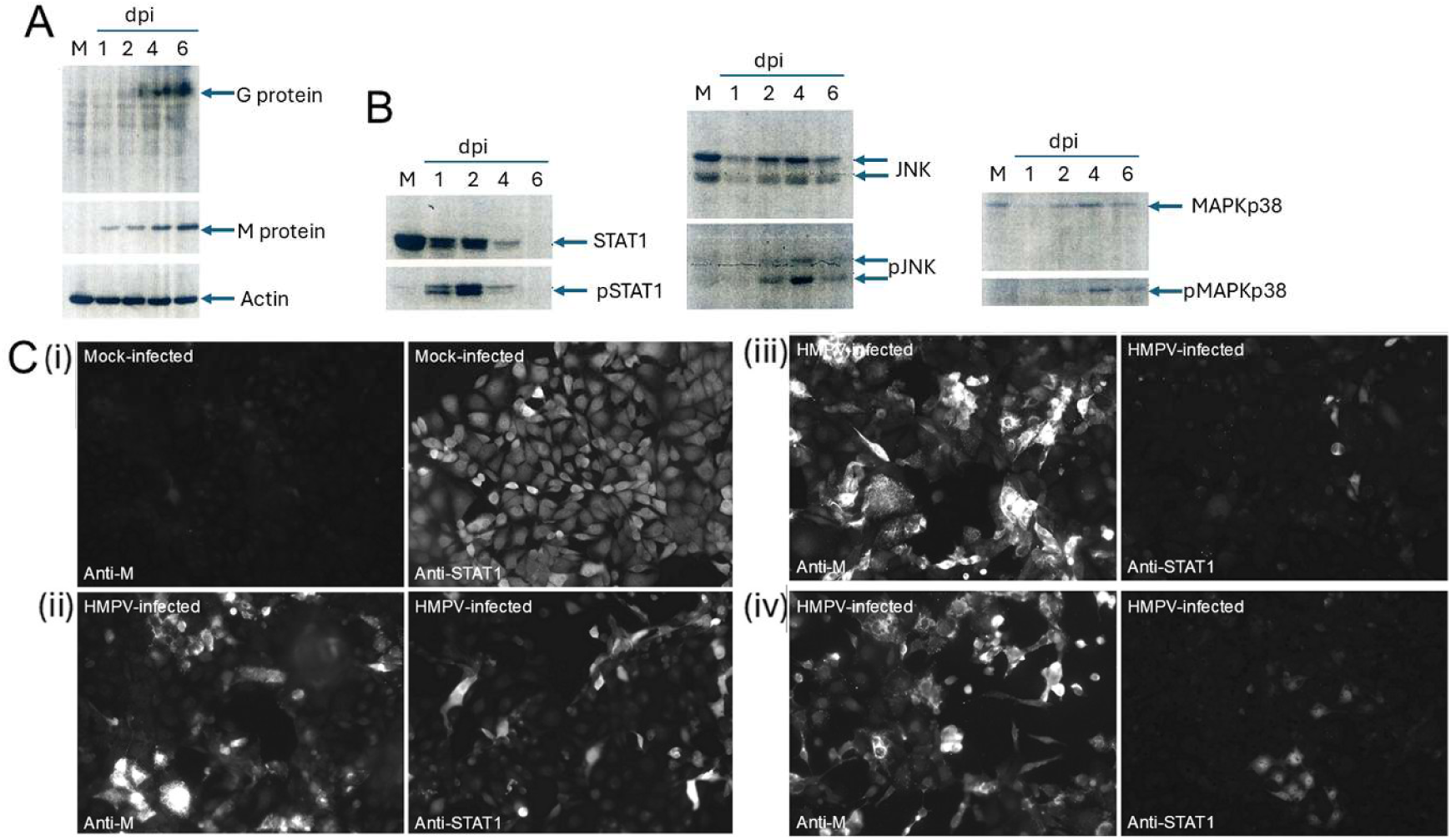
Temporal activation of STAT1, JNK and MAPKp38 signalling pathways in the multiple cycle infection of HMPV in LLC-MK2 cell monolayers. LLC-MK2 cell were mock-infected (M) and infected with HMPV using a moi of 0.01 and at 1 day-post infection (dpi), 2dpi, 4 dpi and 6 dpi cell lysates were prepared. The cell lysates were Western blotted onto PVDF membranes and probed with **(A)** anti-G and anti-M and **(B)** anti-STAT1, anti-pSTAT1 anti-JNK, anti-pJNK anti-MAPKp38 and anti-pMAPKp38 as indicated. The probing with anti-actin is the loading control. In each case protein bands corresponding to the expected size are indicated (black arrow). **(C)** In a parallel analysis duplicate (i) mock-infected monolayers and cell monolayers infected with HMPV at (ii) 2 dpi, (iii) 4 dpi and (iii) 6 dpi were stained with either anti-M or anti-STAT1 and imaged by immunofluorescence microscopy (objective x20 Magnification). The mock-infected cell monolayer pair was stained at an equivalent of 6dpi.

The reduced levels of the STAT1 protein following HMPV infection was confirmed by using IF microscopy to examine infected cells stained using anti-STAT1 and anti-M. The LLC-MK2 cell monolayers were mock-infected and infected with HMPV using a moi of 0.01 and at between 2, 4 and 6 dpi the cells were stained with either anti-M or anti-STAT1 and imaged using IF microscopy (Fig. 9C). In the mock-infected cells no anti-M staining was detected but widespread anti-STAT1 staining across the whole cell monolayer was observed (Fig. 9C(i)). This was consistent with the presence of endogenous basal levels of the STAT-1 protein being expressed in these cells. Increased anti-M staining was observed in the HMPV-infected cells at 2, 4 and 6 dpi was consistent with increased virus replication and the spread of virus infection in the cell monolayer (Fig. 9C (ii) to (iv)). In contrast, there was a concomitant decrease in the levels of anti-STAT1 staining from 2 to 6 dpi (Fig. 9C (ii) to (iv)), and the imaging data was consistent with the immunoblotting data described above. These data collectively suggested that HMPV infection led to reduced STAT1 expression in the LLC-MK2 cells, which as a consequence would be expected to impair interferon signalling as the infection proceeded across the cell monolayer at the later stages of infection. Given the low moi used in the assay, it is currently not clear if during the initial stages of the HMPV infection trans-acting factors are induced in the initial HMPV-infected cells that then down-regulate STAT1 expression in other surrounding cells.

### 3.4 Conclusion

In this study we show that under condition of low moi the HMPV transmits on cell monolayers via two distinct pathways, and in addition two distinct HMPV virus particle morphologies appear to be involved. The short-range transmission occurs via cell-associated virus filaments, and a longer-range transmission that involves cell-free virus. The virus filaments were the main structure detected on the HMPV-infected cells that were cell-associated, but on the distal ends of these structures we also observed that distinct spherical structures (i.e. VPRM) were formed. The spherical structures exhibited a significantly larger width than the width of the virus filaments and their increased staining intensity at the distal ends of the virus filaments suggested an accumulation of the virus glycoproteins at this location. We noted that a proportion of these spherical structures appeared to be detached from the virus filaments and that appear to be directly associated with the adjacent non-infected cells in the cell monolayer. This suggests that the extension of the virus filaments to the non-infected cells may allow the attachment of the spherical structures to the surface of cells that are adjacent to the initially infected cell. Our data also suggests that the spherical particles may be a primary source of the cell-free virus infectivity that we detected later in infection and whose appearance correlated with the spread of infection across the whole cell monolayer. This suggestion is supported by previous studies which have demonstrated the presence of spherical-shaped cell-free HMPV particles in HMPV preparations using transmission electron microscopy [1]. This is similar in essence to the transmission of RSV on permissive cell monolayers reported previously [16], and in this context a model of virus transmission involving changes in virus particle morphology has recently been proposed for the RSV [18]. We have previously also reported increased levels of F protein at the distal ends of the virus filaments that form on RSV-infected cells when infected with RSV using a low moi [25]. This suggests that the increased distribution of the respective F protein at the distal ends of the virus filaments may be a common feature in cells infected with RSV and HMPV. This facet of the virus assembly process of the HMPV and RSV viruses may be functionally relevant during the process of efficient virus transmission. Although in the previous study of RSV there was no direct evidence for a spherical particle morphology on the distal ends of the virus filaments that form on RSV-infected cells, detached spherical RSV structures on the surface of cells were also detected later in infection but their provenance remained unknown at that time.

Previous studies employing the recombinant expressed HMPV F and G proteins in LLC-MK2 cells led to the formation of filamentous virus-like particles (VLPs) on the surface of these cells. This suggested that trafficking signals in one or other of the virus glycoproteins existed that trafficked these proteins into filamentous projections that contained F-actin and lipid rafts that serve as the sites of HMPV particle assembly [6]. However, in these previous studies the formation of spherical structures on the distal ends of the filamentous VLPs were not apparent, suggesting that the formation of the spherical particle morphology is a consequence of the HMPV infection in these cells. This further suggests that virus-induced changes in cell physiology may be involved in the formation of the spherical particle morphology. For example, signalling pathways may be activated during HMPV infection but are not activated in the non-infected cells expressing the F and G proteins. In this context activation of JNK and MAPKp38 pathways was observed in the HMPV-infected cells, which coincided with the time of increased virus replication. In cells infected with the closely related RSV activation of the JNK and MAPKp38 pathways is associated with increased virus replication [23,24]. It is currently unclear if the activation of specific signalling pathways such as the JNK and MAPKp38 networks are required for the formation of the spherical virus structures and this will require further examination. Furthermore, if these signaling pathways do play a role in facilitating the release of cell-free HMPV infectivity it is unclear if this is part of a specific mechanism that leads to virus particle release or if this occurs via a non-specific mechanism e.g. mechanical detachment of the virus due to virus-induced cell membrane damage. The biological significance of the activation of these signaling pathways during HMPV replication and the detailed mechanism behind the formation of the cell-free virus infectivity will require further investigation to address these questions.

Our data suggests that the spherical particles are formed from the virus filaments, which further suggests that the formation of the virus filaments maybe a prerequisite for the formation of the spherical virus particle. This further suggests that virus filament formation may be a prerequisite for the increased levels of the cell-free virus infectivity and spread of infection. In this context evidence for a role of the Rho GTPases Cdc42, Rac1 and RhoA in HMPV virus filament formation has been reported [14]. This is similar to the situation for RSV were activation of the Rac1 and RhoA proteins in RSV-infected cells mediate virus filament formation that facilitate virus transmission [15]. Interestingly prenylation of Rac1 and RhoA rho GTPases is required for trafficking of these proteins to the cell membrane during RSV replication, and in this context the cells mevalonate metabolic pathway provides lipid moieties for protein prenylation. Inhibiting the activity of the key enzyme HMGCR (3-hydroxy-3-methylglutaryl-coenzyme A reductase) in the mevalonate metabolic pathway by using statin-based drugs also prevents protein prenylation and inhibits RSV filament formation and virus transmission via an actin-dependent mechanism [26–28]. The involvement of rho GTPases in HMPV filament formation suggests that similar drugs may also be effective in inhibiting HMPV filament formation and virus transmission, although this possibility needs further investigation.

## Acknowledgments

We thank the National Medical Research Council of Singapore for research funding. We thank Geoff Toms and Fiona Fenwick for providing the HMPV F and G antibodies and the HMPV isolate, and Raihan Jumat and Liat Hui Loo for technical assistance.

## References

1. van den Hoogen, B.G, de Jon, JC., Groen, J, Kuiken, T., de Groot, R., Fouchier, R.A.M, and Osterhaus, ADME. (2001) A newly discovered human pneumovirus isolated from young children with respiratory tract disease Nature Medicine 7, 719–724

2. Ngwoke, I., Ahmed, M.M., Gideon, J.A., Olalekan, J.O., Danladi, N.P., Agboola, A.O., Abdullahi, Y.B., Oso, T.A., Adebayo, U.O., Eshun, G. and Lucero-Prisno, D.L. (2026). Molecular characteristics, epidemiological trends, and public health implications of human metapneumovirus (hMPV): a review (2026). Virology 619: 110897

3. Date, K., Antoniou, E., Polkowska-Kramek, A. et al. (2026). The Global Burden of Human Metapneumovirus in High-Risk Adults: A Systematic Literature Review and Meta-Analysis. J Epidemiol Glob Health (2026). 10.1007/s44197-026-00567-2

4. Schowalter, R.M., Smith, S.E. and Dutch, R.E. (2006). Characterization of human metapneumovirus F protein-promoted membrane fusion: critical roles for proteolytic processing and low pH. J Virol 2006, 80:10931–10941.

5. Liu, L., Bastien, N. and Li, Y. (2007) Intracellular processing, glycosylation, and cell surface expression of human metapneumovirus attachment glycoprotein. J Virol 2007, 81:13435–13443.

6. Loo, L.H, Jumat, M.R., Fu, Y., Ayi, T.C., Wong, P.S., Tee, N.W.S., Tan, B.H. and Sugrue, R.J. (2013). Evidence for the interaction of the human metapneumovirus G and F proteins during virus-like particle formation. Virol J 10, 294

7. Low K.W., Tan T., Ng K., Tan B.H. and Sugrue RJ. (2008) The RSV F and G glycoproteins interact to form a complex on the surface of infected cells. Biochem Biophys Res Commun. 366, 308–13.

8. Huong, T.N., Lee, Z.Q., Lai, S.K., Lee, H.Y., Tan, B.H. and Sugrue, R.J. (2024) Evidence that an interaction between the respiratory syncytial virus F and G proteins at the distal ends of virus filaments mediates efficient multiple cycle infection Virology 591:109985.

9. McGinnes Cullen, L., Luo, B., Wen, Z., Zhang, L., Durr, E. and Morrison, T.G. The respiratory syncytial virus (RSV) G protein enhances the immune responses to the RSV F protein in an enveloped virus-like particle vaccine candidate J. Virol., 97 (1) (2023), Article e0190022

10. van den Hoogen, B.G., Bestebroer, T.M., Osterhaus, A.D. and Fouchier, R.A. (2002) Analysis of the genomic sequence of a human metapneumovirus. Virology 2002, 295:119– 132.

11. Biacchesi, S., Skiadopoulos, M.H., Boivin, G., Hanson, C.T., Murphy, B.R., Collins, P.L. and Buchholz, U.J. (2003). Genetic diversity between human metapneumovirus subgroups. Virology, 315:1–9.

12. van den Hoogen B.G., Herfst, S., Sprong, L., Cane, P.A., Forleo-Neto, E., de Swart, R.L., Osterhaus, A.D. and Fouchier, R.A. (2004). Antigenic and genetic variability of human metapneumoviruses. Emerg Infect Dis, 10:658–666.

13. Jumat, M.R., Huong, T.N., Wong, P., Loo, L.H., Tan, B.H., Fenwick, F., Toms, G.L. and Sugrue, R.J. (2014) Imaging analysis of humanmetapneumovirus-infected cells provides evidence for the involvement of F-actin and the raft-lipid microdomains in virus morphogenesis. Virology Journal 2014 11:198. Virol J 2013 Sep 25:10:294.

14. El Najjar F, Cifuentes-Muñoz N, Chen J, Zhu H, Buchholz UJ, Moncman CL, et al. (2016) Human metapneumovirus Induces Reorganization of the Actin Cytoskeleton for Direct Cell-to-Cell Spread. PLoS Pathog 12(9): e1005922.

15. Sugrue, R.J. and Tan B.H. (2023). Defining the Assembleome of the Respiratory Syncytial Virus. Subcell Biochem. 1062023; 106:227–249

16. Huong, T.N., Iyer Ravi, L., Tan, B.H. and Sugrue, R.J. (2016). Evidence for a biphasic mode of respiratory syncytial virus transmission in permissive HEp2 cell monolayers. Virol J. 13, 12.

17. Huong, T.N., Yan, Y., Jumat, M.R., Lui, J., Tan, B.H., Wang, Y. and Sugrue, R.J. (2018) A sustained antiviral host response in respiratory syncytial virus infected human nasal epithelium does not prevent progeny virus production. Virology. 521:20–32.

18. Sugrue, R.J. and Tan, B.H. (2025). The link between respiratory syncytial virus (RSV) morphogenesis and virus transmission: towards a paradigm for understanding RSV transmission in the upper airway. Virology, 604, 110413.

19. Fenwick, F., Young, B., McGuckin, R., Robinson, M.J., Taha, Y., Taylor, C.E. and Toms, G.L. (2007) Diagnosis of human metapneumovirus by immunofluorescence staining with monoclonal antibodies in the North-East of England. J Clin Virol. ; 40:193–196.

20. Cannon, M.J. (1987) Microplaque immunoperoxidase detection of infectious respiratory syncytial virus in the lungs of infected mice. Journal of virological methods, 16 (4), 293–301.

21. Ingram, R.E., Fenwick, F., McGuckin, R., Tefari, A., Taylor, C. and Toms G.L. (2006). Detection of human metapneumovirus in respiratory secretions by reversetranscriptase polymerase chain reaction, indirect immunofluorescence, and virus isolation in human bronchial epithelial cells. J Med Virol , 78:1223–1231.

22. Duncan CJA, Randall RE, Hambleton S. (2021) Genetic Lesions of Type I Interferon Signalling in Human Antiviral Immunity. Trends Genet. 2021 37(1):46–58.

23. Choi, M.S., Heo, J., Yi, C.M., Ban, J., Lee, N.J., Lee, N.R., Kim, S.W., Kim, N.J. and Inn, K.S. (2016). A novel p38 mitogen activated protein kinase(MAPK) specific inhibitor suppresses respiratory syncytial virus and influenza A virus replication by inhibiting virus-induced p38 MAPK activation. Biochemical and Biophysical Research Communications Volume 477, 311–316

24. Caly, L., Li, H.M., Bogoyevitch, M.A and Jans, D.A. (2017). c-Jun N-terminal kinase activity is required for efficient respiratory syncytial virus production. Biochemical and Biophysical Research Communications. . 483, 64–68.

25. Tra Nguyen Huong, Zhi Qi Lee, Soak Kuan Lai, Hsin Yee Lee, Boon Huan Tan, Richard J. Sugrue, (2024). Evidence that an interaction between the respiratory syncytial virus F and G proteins at the distal ends of virus filaments mediates efficient multiple cycle infection. Virology 591, 109985

26. Gower TL, Peeples ME, Collins PL, Graham BS. (2001). RhoA is activated during respiratory syncytial virus infection. Virology. 10;283(2):188–96.

27. Ravi LI, Liang L, Wong PS, Brown G, Tan BH, Sugrue RJ. (2013) Increased hydroxymethylglutaryl coenzyme A reductase activity during respiratory syncytial virus infection mediates actin dependent inter-cellular virus transmission. Antiviral Res. 100(1):259–268.

28. M Malhi, MJ Norris, W Duan, TJ Moraes and JT Maynes (2021). Statin-mediated disruption of Rho GTPase prenylation and activity inhibits respiratory syncytial virus infection. Commun Biol 4, 1239.

